# Boosting reliability when inferring interactions from time series data in gene regulatory networks

**DOI:** 10.1101/2025.02.17.638617

**Authors:** Mattia Greco, Federico Ricci Tersenghi, Olivier C. Martin

## Abstract

In the context of the dynGENIE3 [13] approach for inferring regulatory network interactions from time-series data, we show that it is possible to modify that algorithm to significantly enhance its prediction reliability. To quantify the level of reliability, we used ground-zero truths based on simulated datasets generated by the GeneNetWeaver [22] tool. Our work introduces novel methods leveraging time-lagged correlations and estimators of mRNA decay rates, leading to significantly improved driver-target inference. Additionally, a temperature-based rescaling of priors was developed to further enhance prediction reliability. Results demonstrate substantial improvements in performance with a particularly notable increase in AUPRC scores. These advances underscore the possible gains resulting from incorporating priors into gene regulatory network inference.

## 1 Introduction

Natural selection has led nearly all biological organisms to tightly regulate the processes important for their survival. At the heart of those control mechanisms lie so called gene regulatory networks (GRNs) [10]. At a coarse level of description, a GRN can be summarized as list of driver-target gene pairs, while a more detailed description might include the rules for the expression of the target genes as a function of the state of their drivers, requiring for instance a dynamical systems framework. Since experimental validation of even a single putative driver-target pair requires a major effort, much theoretical work has been put into developing ways to infer such interactions from high-throughput gene expression data [4], [5], [15]. In particular, RNA-seq [25] is now a cheap way to quantify the abundances of transcripts and of providing associated genome-wide data. Of course, RNA abundances inform only on part of the dynamics of genes (the protein level is considered to be more important for function, but quantifying protein levels remains a challenge to date). Nevertheless, RNA-seq measurements are now sufficiently accessible to become the entry point for inferring regulatory interactions. As a result, numerous computational tools have been published for inferring putative interactions between genes from such data. Initially, most efforts were based solely on “co-expression” [26] so that only non oriented pairs were proposed, in other words it was not clear which gene of a pair was the driver vs the target. To lift that ambiguity, methodologies switched to frameworks that infer the “rule” for going from the expression level of a list of driver genes to the expression level of a given target gene. The hope therein is to identify the true causal factors controlling each target gene. Such a program is particularly promising when the biological system is time dependent, for instance when it responds to a perturbation or undergoes a spontaneous developmental process. Thus in the present work we focus on the question of inferring oriented interactions in GRNs based on so-called “longitudinal” data, that is data taken at a succession of different time points.

The community of researchers working on this objective has been structured by annual competitions in which any author could provide the code of their algorithm that would then be blindly benchmarked on test examples. These series of DREAM (Dialogue for Reverse Engineering Assessments and Methods) [23] competitions lasted for a half a dozen years, from which it transpired that the winners were generally based on using Random Forests to “learn” the rules between drivers and targets. In terms of implementation, the dynGENIE3 algorithm [13] came out as the winner several times for the reliability of its predictions, in addition to being particularly fast in terms of execution time and being able to integrate both longitudinal data and more standard co-expression-type data. The motivation of the present work is to investigate to what extent the Random Forest approach can be enhanced by introducing priors when constructing the underlying trees. For convenience, we do so by modifying the dynGENIE3 (open source) algorithm as it also is the most widely used software for studying GRNs with Random Forest.

We will first cover the heart of the DREAM benchmarking methodology in which one assigns a performance measure to an algorithm based on its AUROC and AUPRC values. These quantities are standard measures in the community of binary classifiers that in our context correspond to assigning interactions to be either true or false. To obtain such performance measures it is thus necessary to know the “ground truth”, that is which interactions are indeed the correct ones. Because biological knowledge is still incomplete even for very well studied organisms, in practice one has no choice but to use simulated data for providing ground truths. The DREAM competitions provided a standard framework for generating such data via the software GeneNetWeaver which allows quite realistic regulatory rules. We thus follow this methodology throughout this paper. Given that the challenge of inferring the drivers of a target gene likely becomes more difficult as its number of drivers increases, we have systematically controlled for that number, successively considering graphs with in-degrees of 1, 2, 3 and 4.

In the following parts of this paper, we tackle the task of modifying dynGENIE3 to improve its performance. In section 3 we start by defining time correlation functions. We then quantify in sub-section 3.1 the reliability of using these to predict the sign of interactions. The main part of our work is presented in sub-section 3.3 where we show that the use of time-lagged correlations for introducing priors and for estimating messenger RNA decay rates can significantly improve the reliability of the Random Forest inference of interactions. Lastly, sub-section 3.4 provides a way to fine-tune the priors based on a non linear transformation motivated by a temperature parameter. Our conclusions are presented in the last section.

## 2 Methods

### 2.1 Data generation and benchmarking

Many different tools have been developed to infer interactions in gene regulatory networks [18], [11], [14], [12], [3], [16], and it is important to assess the reliability of their predictions. This task was realized by the DREAM (Dialogue for Reverse Engineering Assessments and Methods) [23] competitions via standardized benchmarking measures of network inference performance. In the series of DREAM competitions, participants were given data (either steady state or time series) generated by simulating transcriptional dynamics, from which they had to reconstruct the underlying network that had been used to generate the dynamics. For the present study, we will focus on the dynGENIE3 [13] algorithm which was the winner of the DREAM4 and DREAM5 competitions. The simulator that was developed to generate the data for these competitions is called GeneNetWeaver (GNW) [22]. GNW^1^ must first be given the topology of a network with associated parameters (for example, number of nodes, in-out degrees, noise levels, etc.). Then, using a quite realistic model, GNW simulates the dynamics of transcription and translation of the genes in the network. The choices made for the dynamical equations are motivated physically and biologically in [1]. The supplementary material of [22] provides more detailed explanations. Since we will use GNW to produce simulated data, we briefly describe the underling procedure here.

The GNW framework allows both additive (independent) and multiplicative (synergistic) interactions. For a given gene *i*, it assumes the following set of coupled ODEs:

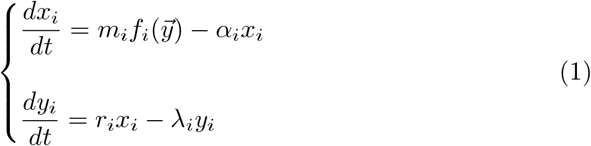

in which *x*_*i*_ is the associated mRNA concentration, *y*_*i*_ the protein concentration, *α*_*i*_ and *λ*_*i*_ are the decay rates of these mRNAs and proteins, respectively, and *m*_*i*_ and *r*_*i*_ are the maximum transcription and translation rates. The function *f*_*i*_ is called the activation function of the gene *i* and depends on the concentrations of the proteins that control gene *i*’s transcription. The form of the function used to generate the data is fairly complicated and does not concern us here since we are mainly interested in just inferring which are the driver genes.

The integration of these equations provides noiseless trajectories 1 (bottom left) for the expression levels in time. In general, these processes (transcription and translation) are affected by intrinsic noise [9] (fluctuations at the molecular level) and experimental measuring errors. To introduce stochasticity into the data, two Gaussian white noises can be added to each equation: one that depends on the “production” (first terms of eqs. 1) and another depending on the “degradation” term (second terms of eqs. 1). The resulting stochastic equations are integrated à la Stratonovich 1 (bottom right). Finally it is possible to add an extra noise to simulate experimental errors. Figure 1 illustrates these steps of the process of time series data generation.

**Figure 1:**
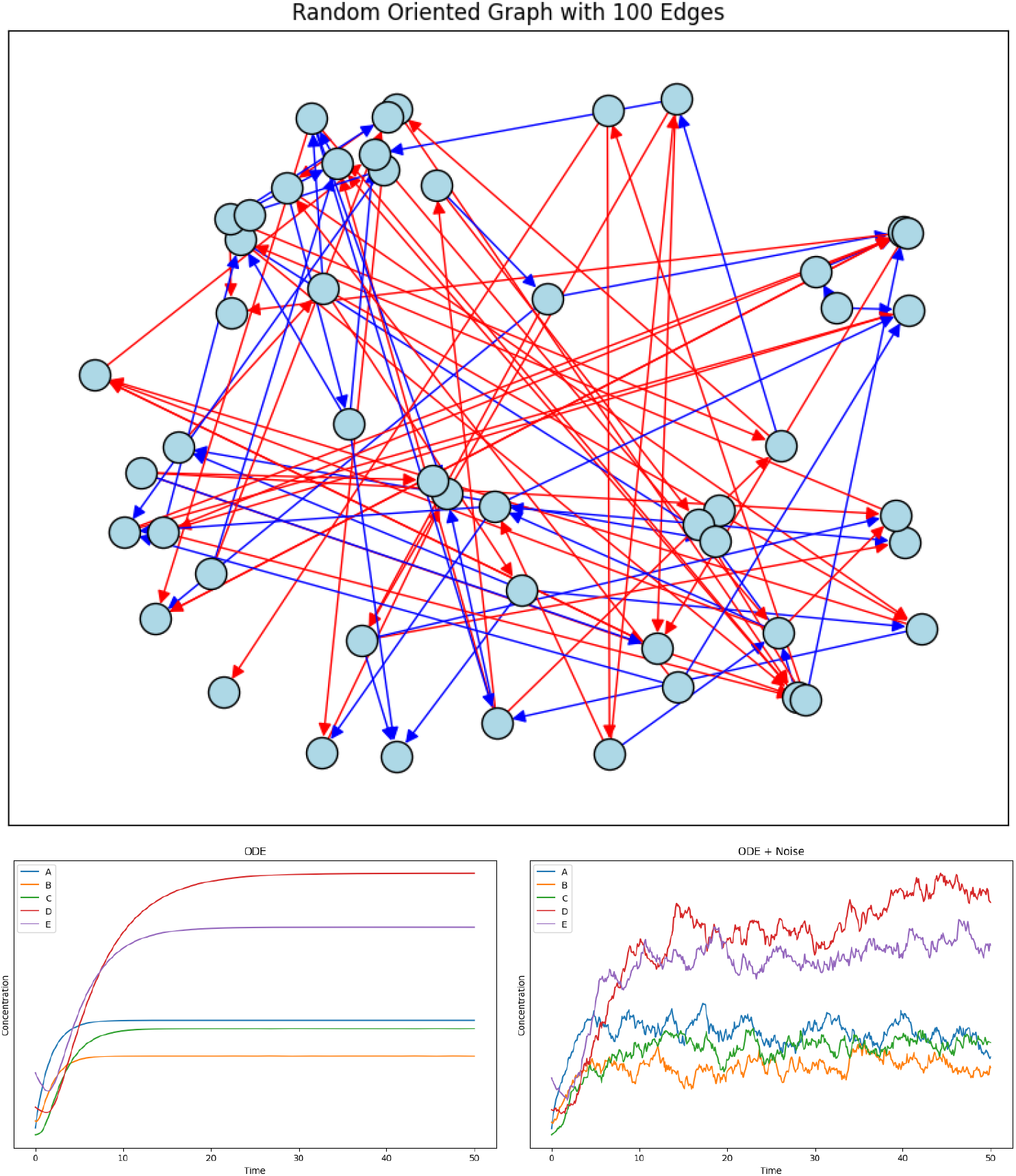
The process of generation of *in silico* time series. Top: Example of a graph, namely a list of nodes (genes) and oriented edges (interactions). Bottom left: Simulation generating the noiseless time-evolution for some of the genes in the network. Bottom right: Noisy realization of the same dynamical model.

In the following, we will be concerned with the decay rates of messenger RNAs (the *α*_*i*_s of eq.1) and the nature of the interactions (i.e., inhibitory or activatory). The ground-truth values of these parameters, used by GNW to generate the data, can be compared to inferred values, thereby allowing one to measure the quality of an algorithm’s inference.

The datasets we will be working with are composed of ensembles of random graphs having 100 genes. Specifically, we will consider random graphs having a fixed in and out degree. Our values for the in and out degrees *d* will be set to 1, 2, 3 and 4. A gene’s in degree refers to the number of incoming interactions while its out degree is its number of outgoing interactions. To illustrate this, we display some corresponding graphs in Fig.2 (but having only 10 genes for ease of visualization). The total number of (oriented) interactions *N*_*I*_ in the network satisfies:

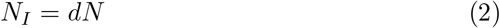

where N is the number of genes. We will need to infer 100, 200, 300 and 400 interactions out of the 10000 possible ones depending on the in-out degree. Of course these networks are not completely realistic since the in-out degree of the nodes in biological gene networks are not fixed and can even exhibit scale invariance properties [4]. However what we are able to see, by fixing the inout degree, is how the algorithm’s reliability changes with the average node degree.

**Figure 2:**
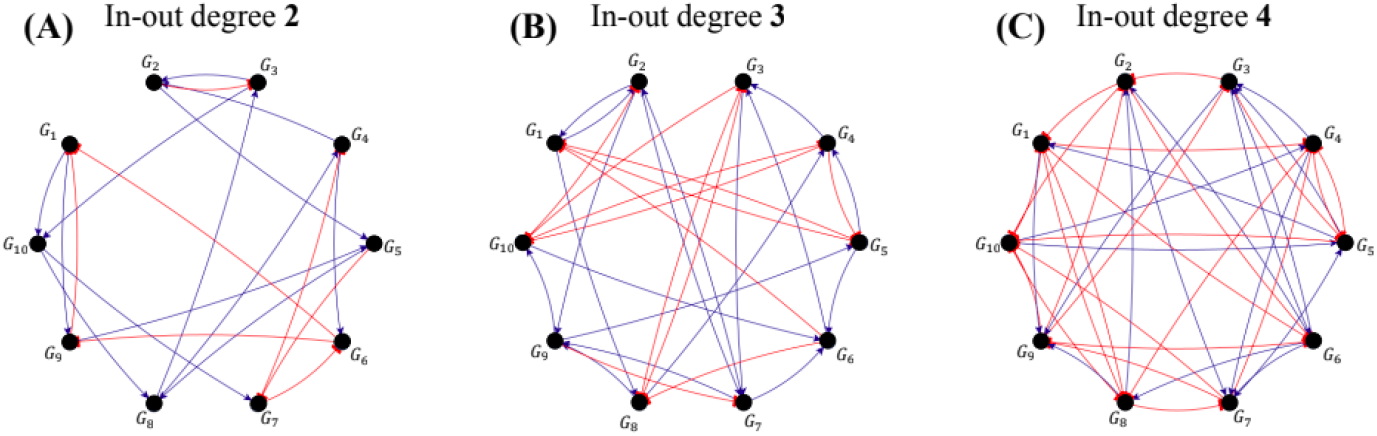
Examples of networks having 10 genes, with in and out degrees of 2, 3 and 4.

**Figure 3:**
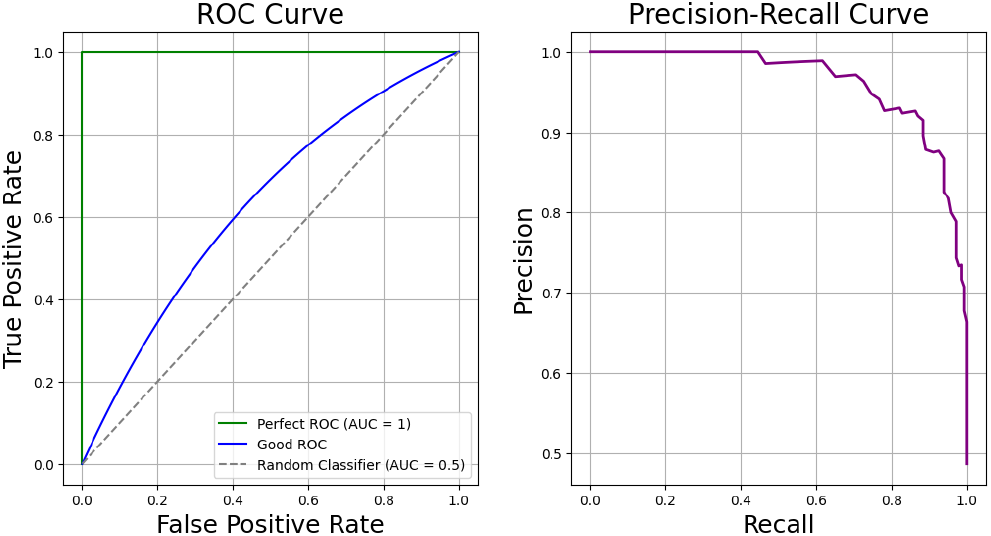
Examples of receiving operating curve and precision-recall curve.

Given such networks, our benchmarking datasets are created via GNW and following standard choices of the DREAM competitions. Specifically, GNW simulates the gene expression dynamics produced by ordinary differential equations, leading to 10 time series consisting of 21 points equally spaced in time.

### 2.2 Metrics for quantifying reliability

The metrics generally chosen to measure the performance (reliability) of GRN inference algorithms are fairly standard. In particular, we focus on the Area Under the Receiving Operating Curve (AUROC) and the Area Under the Precision Recall Curve (AUPRC) [17]. These two quantities are very commonly used in general classification tasks and are directly applicable here when predicting which edges are present in the ground truth GRN used by GNW for generating time series.

The receiving operating curve for a binary classificator compares the True Positive Rate (TPR) against the False Positive Rate (FPR) 3. Clearly, if the TPR and the FPR are equal, the classifier does no better than random, while in the ideal scenario the FPR is 0 and the TPR is 1. The area under this curve generally takes values between 0.5 (no predictive power) and 1 (perfect classifier). On the other hand, the precision recall curve 3 compares the precision of the algorithm, defined as:

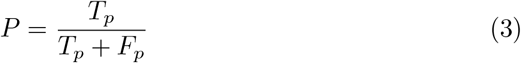

where *T*_*p*_ is the number of true positives and *F*_*p*_ is the number of false positives, against the recall, defined as:

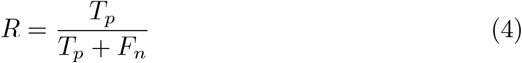

in which *F*_*n*_ is the number of false negatives. So, the curve in 3 will initially be flat, staying at its maximum of 1 as long as the prediction of positives has no errors. The area under the precision recall curve will be a number between 0 (worst scenario) and 1 (perfect scenario). In the case concerning us here (predicting interactions in GRNs using dynGENIE3 and improvements thereof), the output of the algorithm is a matrix containing the “importance” of each putative interaction. These importances are between 0 and 1. One could choose a threshold for these importances, setting to 0 all the importances below that threshold and setting to 1 all the ones above, leading to a prediction for which edges are to be considered as true. Once this is done, the obtained network could be compared to the ground truth to estimate the number of true and false positives. To avoid choosing an arbitrary threshold, the procedure used in the DREAM competitions consists in averaging over all possible thresholds; it is easy to see that this corresponds to determining the AUROC or AUPRC values.

### 2.3 dynGENIE3

The algorithm we focus on is called dynGENIE3 [13]. dynGENIE3 was the winner of the DREAM4 and DREAM5 competitions, which justifies our use of it. Furthermore, it is extremely fast, and allows analyzing steady-state and time series data jointly.

The algorithm is based on a random forest regressor [7]. Given *N* genes, we represent a time series data set of *n* points in time as:

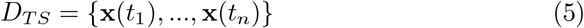

where **x**(*t*_*j*_) ∈ **R**^*N*^ is a vector containing the expression levels of all *N* genes at time *t*_*j*_. In the same way we can include steady-state data as:

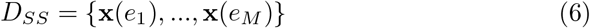

where we denote by *e*_*k*_ the experimental condition *k* which is assumed to be some equilibrium state of our system. dynGENIE3 models the expression level of gene *i* with the following ODE:

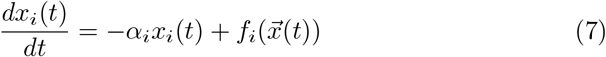

in which *α*_*i*_ is the decay rate of the messenger RNAs for that gene. Note that the transcription rate of this gene *i* is given by an arbitrarily complicated function *f*_*i*_ whose argument 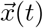 consists of the expression levels of the genes driving it. This equation is discretized as:

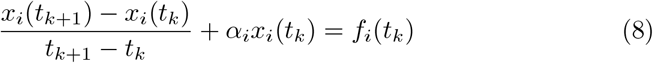

for *k* = 1, …, *N* − 1. From this discretization the learning set for the regressor is built:

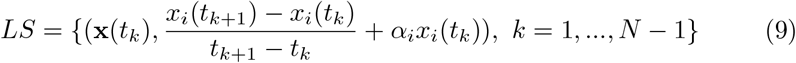

from which it is possible to fit the function *f*_*i*_. If steady-state data are also present, the learning set is augmented by expression levels where the time derivative is zero. Clearly, once this minor modification is made, one can use both types of data together.

It is clear that dynGENIE3 does not explicitly consider the protein step of equation 1 in its modeling. Ignoring the protein step makes the modeling “effective” and thus the *α*_*i*_ parameters of equation 1 are not directly related to the ones of equation 7. This fact matters since the decay rates are inputs to the algorithm and its performance depends on them as highlighted already in [13].

After creating this learning set, a regression tree ensemble fits the *f*_*i*_. Regression trees [7] [8] use binary tests to split the data based on single inputs in order to minimize the variance of the output. The threshold of each possible split is determined while the tree grows. At each node of the tree *K* possible drivers for the given target are extracted at random and are tested in order to determine the best split. In the original version of the algorithm these *K* drivers are randomly drawn from all possible drivers.

In this paper we are going to show how the performance of dynGENIE3 can benefit from introducing prior probabilities in the drawing of the genes. The predictions for each tree are then averaged over all trees. Such an averaging process usually improves significantly the predictive power and is typically referred to as the “wisdom of crowds” [24]. Given the tree regressor, the network is inferred by assessing the importance that each input has in predicting the output. The score of each interaction is calculated as the mean impurity decrease measure [8] which is defined as:

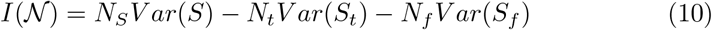

where *S* is the set of samples that reach node 𝒩 while *N*_*S*_ denotes the cardinality of the set *S. S*_*t*_ (*S*_*f*_) denotes the subset for which the test is true (false) and *N*_*t*_ (*N*_*f*_) its cardinality. Given *I* for a single tree, the weight *w* of a variable is taken as the average of Eq. 10 over all the splits in which that variable is used. Clearly if a variable is not selected at all its importance will be 0, while if it is selected close to the root of the tree its importance will, typically, be high.

The sum of the importances over all the inputs for a given gene, i.e., the sum over all possible drivers of a given target, is usually close to the total variance of the output if the tree is deep:

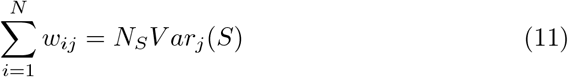

where *V ar*_*j*_(*S*) is the variance of the target gene *j* estimated from *S*. If the scores were used naively, there would be a positive bias for the genes with bigger variations. To avoid any biases, standard practice is to normalize the weights as:

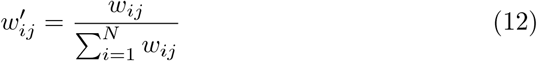

and setting self interactions to 0, i.e., putting *w*_*ii*_ = 0. These weights are then averaged over all trees of the random forest, thereby defining the so called “importances”.

## 3 Results

### 3.1 Equal-time and time-lagged correlations

In this section we introduce correlations between drivers and targets using equal-time or time-lagged values. We denote by *x*_*i*_(*t*_*n*_) the centered expression level of driver *i* at time *t*_*n*_, that is after subtraction of its time-averaged value. We similarly denote by *y*_*j*_(*t*_*n*_) the corresponding value of target *j*. For ease of notation, we define the associated vectors *X*_*i*_ = (*x*_*i*_(*t*_1_), *x*_*i*_(*t*_2_), …, *x*_*i*_(*t*_*N*_)) and *Y*_*j*_ = (*y*_*j*_(*t*_1_), *y*_*j*_(*t*_2_), …, *y*_*j*_(*t*_*N*_)). We can then consider the matrix of (equal-time) linear correlation coefficients:

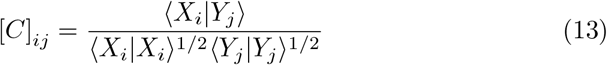

where ⟨−|−⟩ refers to the standard scalar product.

We are also interested in time-lagged correlations that tell us how much the concentration of the *j*-th target at any time is correlated with the concentration of the *i*-th driver at the previous time step. For that we introduce the vectors: 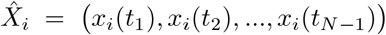 on the one hand and 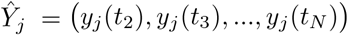 on the other. From these we define the matrix of time-lagged linear correlation coefficients:

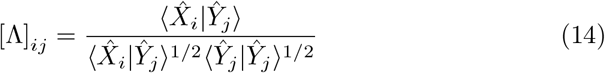

These correlations provide more direct evidence about the putative influence of a driver on a target.

Consider now the distributions of these correlations when using simulated data as generated following the procedures in Methods. The two cases ([*C*] and [Λ]) are displayed in figure 4 for a single random graph of in-degree and out-degree set to 2 for all 100 genes.

**Figure 4:**
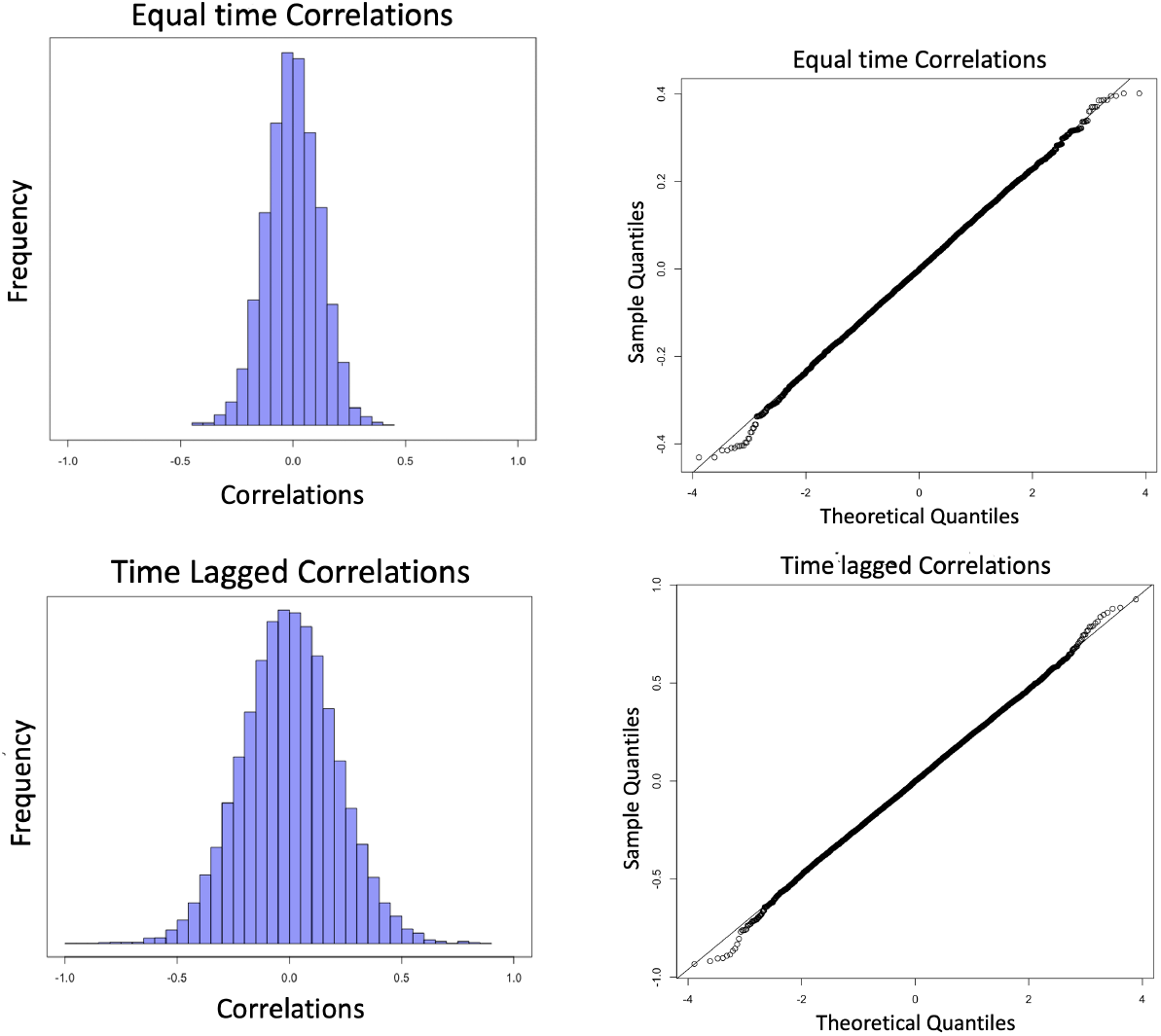
Distribution of equal-time and time-lagged correlations for a random graph having in and out degree of 2 for all genes (top and bottom). On the right the figures display the respective QQ plots, showing that the distributions are close to Gaussian.

For these data we provide the associated QQ plots 4 that allow one to visualize deviations from Gaussian distributions. Clearly these distributions are very close to Gaussian. To measure the gaussianity we use the Shapiro-Wilk test on a smaller sample of data^2^. On a resampling of our distribution the test gives a statistics of 0.999 in both cases and p-values of 0.310 for equal-time correlations and of 0.168 for time-lagged correlations. Since the null hypothesis of the test is the normality of the distributions we do not reject it when using this resampling. This allows us to conclude that both distributions are very close to being Gaussian.

We will use that property later but for the moment note that the dispersion of the time-lagged correlations is significantly greater than that of the equal time correlations, suggesting that [Λ] provides more information than [*C*]. To provide further evidence in favor of using time-lagged correlations, we subset these matrices to the pairs (*ij*) that correspond to the true interactions in these simulated data. Averaging over 100 graphs as displayed in Fig.5, we see that the correlations for the real interactions are much more spread out for the time-lagged case, in line with the expectation that these provide more information on interactions than the equal-time ones.

**Figure 5:**
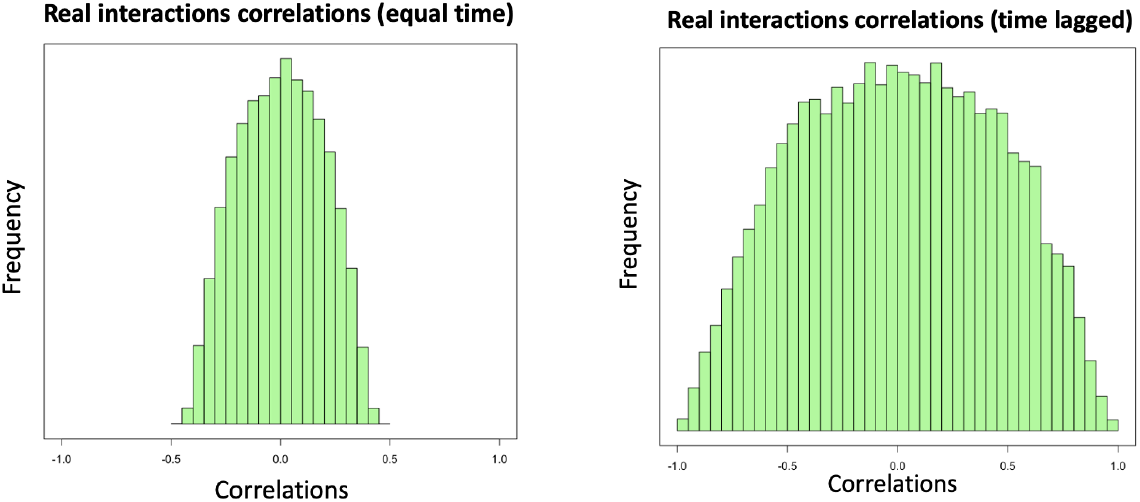
Distributions of correlations when using the real interactions for 100 random graphs (same characteristics as for Fig. 4). Left: equal-time, right: time-lagged.

### 3.2 Predicting the signs of interactions

In a gene regulatory network, when considering a given driver-target pair, the target will typically either activate or repress the expression of the target^3^ [2]. In this section we focus on inferring – again on simulated data – the sign of an interaction (+ for activating and - for repressing) in a GRN using the correlations from Eqs. 13 or 14.

Intuitively, if a driver activates (represses) the expression of a target, we can expect their correlation to be positive (negative). We thus propose to infer the sign of an interaction by taking the sign of either the associated equal-time or time-delayed correlation coefficients. We now quantify the reliability of such an inference on simulated data. For that, we introduce the quantities *r*_+_ and *r*_*−*_ as follows. *r*_+_ is the average over 100 simulated GRNs of the probability of correctly identifying the sign of a positive interaction, while *r*_*−*_ is the analogous quantity for a negative interaction. In both cases, the predicted sign of an interaction is that of the corresponding correlation. In table 1 we can see that such a procedure identifies the signs of an interaction quite accurately. We also see that as the in-out degree increases the retrieval rate decreases.

**Table 1:** Sign retrieval rates for different in-out degrees using different types of correlations.

| Correlations | $r_+$ | $r_-$ | In-out degree |
| --- | --- | --- | --- |
| Time-lagged | $0.96 \pm 0.03$ | $0.95 \pm 0.03$ | 1 |
| Equal-time | $0.94 \pm 0.03$ | $0.93 \pm 0.04$ | |
| Time-lagged | $0.87 \pm 0.04$ | $0.87 \pm 0.03$ | 2 |
| Equal-time | $0.85 \pm 0.04$ | $0.84 \pm 0.03$ | |
| Time-lagged | $0.80 \pm 0.03$ | $0.80 \pm 0.03$ | 3 |
| Equal-time | $0.78 \pm 0.03$ | $0.77 \pm 0.03$ | |
| Time-lagged | $0.75 \pm 0.03$ | $0.75 \pm 0.03$ | 4 |
| Equal-time | $0.73 \pm 0.03$ | $0.73 \pm 0.03$ | |

Moreover, the performance is slightly better when using time-lagged correlations rather than equal-time correlations. This is in line with the expectation that the use of a time lag probes the dependence of target on driver better than if no time lag is introduced.

To understand what drives the error in such inference of the sign of an interaction, we now examine how the error rate depends on the strength of the correlation. For that we separate the cases with correctly and incorrectly inferred signs for the case of time-lagged correlations. The corresponding distributions are displayed in Fig. 6. Clearly, the two distributions have very different shapes. In particular, the cases where one incorrectly predicts sign come from the region of small correlation values, thus giving a bell shaped distribution around 0 for those incorrectly inferred signs. In contrast it is possible to be more confident in the inference of the signs whenever the correlations have an absolute value that is say larger than 0.2 or 0.5 (depending on the in-out degree). Another feature of these plots is their symmetry, which tells us that it is just as easy (or difficult) to predict the signs of positive and negative interactions, a result that was not a priori obvious since that symmetry is not present at the level of the transcriptional dynamics. Moreover it is worth noting that, as the in-out degree of the graphs increases, the overlap between the two different distributions becomes more and more important. This reflects the decrease in the accuracy of retrieval of the signs.

**Figure 6:**
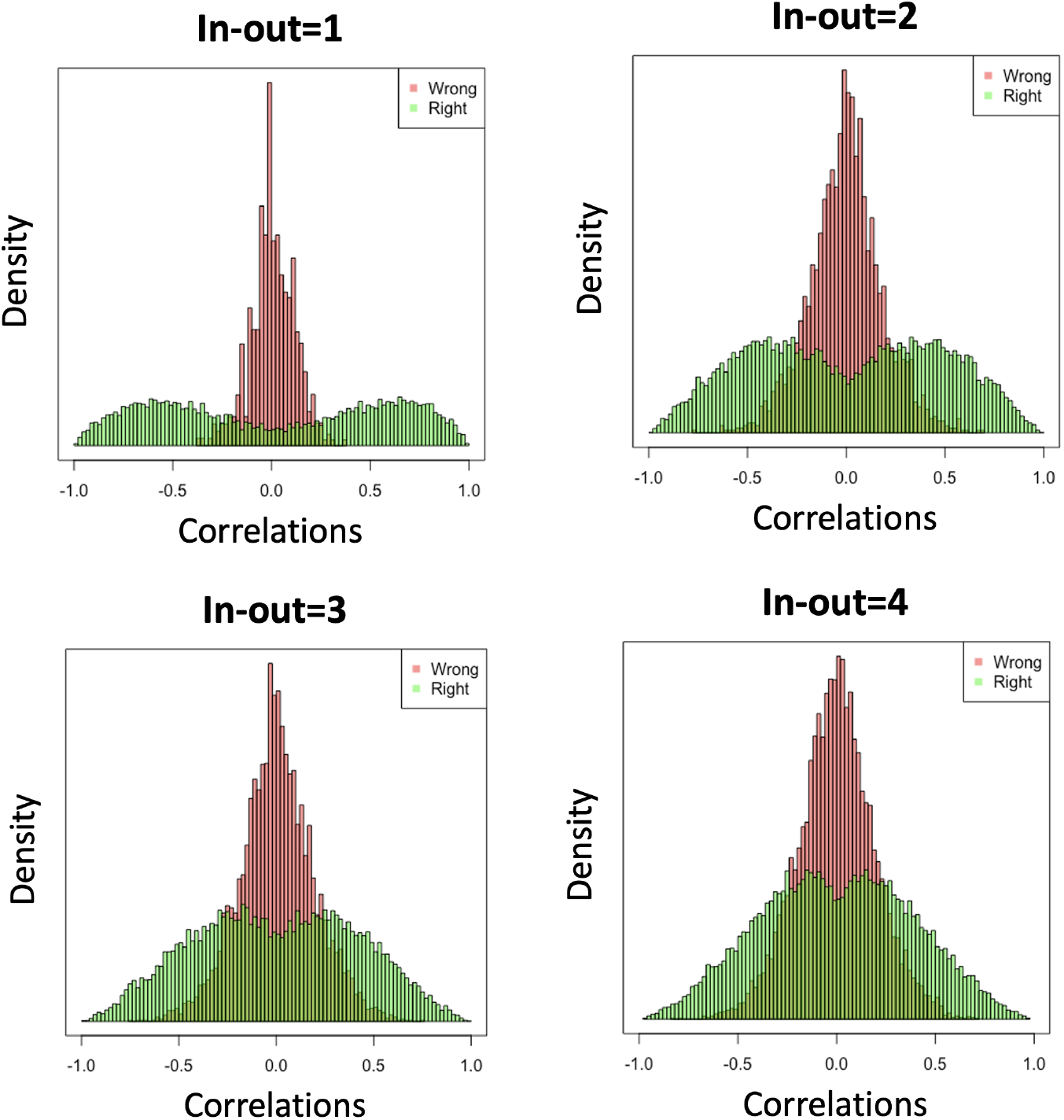
Distribution of time-lagged correlations for correctly (green) and incorrectly (red) identified signs of interactions for different values of the in-out degree.

### 3.3 Priors using time-lagged correlations and modified estimators of mRNA decay rates

In this section we propose two procedures that together significantly enhance the reliability of the dynGENIE3 algorithm.

To ensure full comparability and reproducibility of the results, we fix for the rest of the paper the dynGENIE3 parameters at commonly used values. Specifically, we set 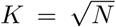 where *K* is the number of genes considered at each node splitting of the trees in the random forest and *N* is the number of genes. Furthermore, we set the number of trees in the random forest to be *n*_*trees*_ = 100 which allows for a fast forest construction. The results we present in the following could likely be further improved by tuning these parameters^4^.

Based on the simulated time-series dataset to analyze, we want to incorporate some priors to improve the GRN inference. Given a target gene, some candidate drivers will show higher correlations with it than others. The first step of our strategy will be to bias the choice of drivers during the construction of trees in the random forest in favor of those putative drivers.

In a random forest, when constructing a regression tree, the algorithm generates split nodes. At each split, *K* drivers are tested and the one that minimizes the variance of the output (leading to the highest information gain) is then selected for the split. In the original dynGENIE3 algorithm, the *K* genes are redrawn at random from the set of *N* genes, for each split node. In our modified approach, we draw the *K* drivers at the beginning of the tree construction so that they are the same for all splits in that tree. Furthermore, they are drawn from a prior distribution dependent on the aforementioned correlations of Eqs. 13 and 14 rather than being drawn uniformly.

In our procedure, the absolute value of these correlation coefficients are taken as priors (up to a normalization constant) in the tree construction step of the random forest. When we do so, we find that the performance of dynGENIE3 increases. This improvement is quantified in Tab. 2 using the AUROC and AUPRC scores, comparing the results of the original dynGENIE3 algorithm with the results from runs using these priors. Although the AUROC scores are only modestly improved, the AUPRC scores are improved quite spectacularly.

We now consider the addition of a second modification to the dynGENIE3 algorithm based on changing that algorithm’s way of estimating mRNA decay rates. The original method for the inference of these parameters, explained in [14], is a bit naive. It takes the maximum and the minimum of the expression levels across the time series and then it fits an exponential between them to find a decay rate. However, it quite often happens that the minimum occurs before the maximum, in which case the inferred “decay” makes no sense. As pointed out by the authors of [14] this method only gives a rough order of magnitude for the decay rates.

Our approach consists in using an inference of the decay rates based on a linear modeling of the dynamics. Specifically, for each gene *i* we model the relaxation to its time-averaged value via a decay rate *α*_*i*_, leading to the simple equation:

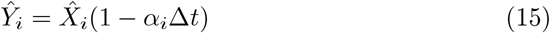

This vectorial equation assumes that the time interval Δ*t* is the same throughout the series, but the equations for the more general case are obvious to write in scalar form. Note that this equation ignores the effects of any interactions^5^. Using Eq. 15 to compute the diagonal elements of the Λ matrix, we find:

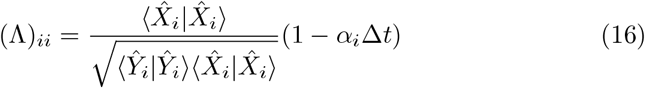

from which one extracts the following estimates of the decay rates:

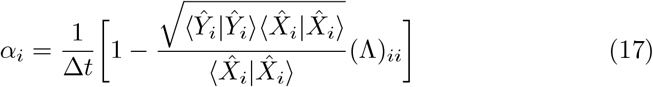

We find that, just as for the default estimates used by dynGENIE3, these decay rates are poorly correlated with the real ones (around 5% correlation in the case of the simulated data produced by GNW). The reason behind this is the fact that we are using an effective model since dynGENIE3 ignores the translation step present in the GNW simulator. Nevertheless, the use of our modified estimates gives particularly good results for the AUPRC, as can be seen in Table 2.

**Table 2:** AUROC and AUPRC for different schemes and in-out degrees.

| Scheme | AUROC | AUPRC | In-out degree |
| --- | --- | --- | --- |
| Original | $0.91 \pm 0.03$ | $0.38 \pm 0.07$ | 1 |
| Priors | $0.91 \pm 0.03$ | $0.56 \pm 0.06$ | |
| Priors+Decay rates | $0.92 \pm 0.03$ | $0.62 \pm 0.03$ | |
| Original | $0.75 \pm 0.02$ | $0.16 \pm 0.03$ | 2 |
| Priors | $0.77 \pm 0.02$ | $0.23 \pm 0.03$ | |
| Priors+Decay rates | $0.78 \pm 0.02$ | $0.30 \pm 0.04$ | |
| Original | $0.66 \pm 0.02$ | $0.11 \pm 0.02$ | 3 |
| Priors | $0.69 \pm 0.02$ | $0.15 \pm 0.02$ | |
| Priors+Decay rates | $0.70 \pm 0.02$ | $0.20 \pm 0.02$ | |
| Original | $0.62 \pm 0.02$ | $0.10 \pm 0.02$ | 4 |
| Priors | $0.64 \pm 0.01$ | $0.13 \pm 0.02$ | |
| Priors+Decay rates | $0.65 \pm 0.01$ | $0.17 \pm 0.02$ | |

**Table 3:** AUROC and AUPRC at different temperatures for different in-out degrees.

| $\beta$ | AUROC | AUPRC | In-out degree |
| --- | --- | --- | --- |
| 0 | $0.89 \pm 0.03$ | $0.32 \pm 0.06$ | 1 |
| 1 | $0.92 \pm 0.03$ | $0.62 \pm 0.06$ | |
| 2 | $0.92 \pm 0.03$ | $0.65 \pm 0.06$ | |
| 3 | $0.92 \pm 0.03$ | $0.71 \pm 0.06$ | |
| 4 | $0.91 \pm 0.03$ | $0.66 \pm 0.06$ | |
| 5 | $0.91 \pm 0.03$ | $0.66 \pm 0.06$ | |
| 0 | $0.74 \pm 0.02$ | $0.14 \pm 0.02$ | 2 |
| 1 | $0.78 \pm 0.02$ | $0.30 \pm 0.04$ | |
| 2 | $0.78 \pm 0.02$ | $0.30 \pm 0.04$ | |
| 3 | $0.78 \pm 0.02$ | $0.36 \pm 0.04$ | |
| 4 | $0.77 \pm 0.02$ | $0.33 \pm 0.04$ | |
| 5 | $0.77 \pm 0.02$ | $0.32 \pm 0.04$ | |
| 0 | $0.70 \pm 0.02$ | $0.17 \pm 0.02$ | 3 |
| 1 | $0.70 \pm 0.02$ | $0.20 \pm 0.02$ | |
| 2 | $0.70 \pm 0.02$ | $0.22 \pm 0.03$ | |
| 3 | $0.70 \pm 0.02$ | $0.22 \pm 0.03$ | |
| 4 | $0.69 \pm 0.02$ | $0.22 \pm 0.03$ | |
| 5 | $0.69 \pm 0.02$ | $0.22 \pm 0.03$ | |
| 0 | $0.65 \pm 0.02$ | $0.14 \pm 0.02$ | 4 |
| 1 | $0.65 \pm 0.01$ | $0.16 \pm 0.02$ | |
| 2 | $0.65 \pm 0.01$ | $0.18 \pm 0.02$ | |
| 3 | $0.65 \pm 0.02$ | $0.18 \pm 0.03$ | |
| 4 | $0.64 \pm 0.01$ | $0.18 \pm 0.02$ | |
| 5 | $0.64 \pm 0.01$ | $0.17 \pm 0.02$ | |

To illustrate this in greater detail, Fig. 7 displays the PR curves (averaged over 100 graphs) for the three different algorithms, when the graphs’ in-out degrees are 1, 2, 3 or 4. We see that whenever we introduce priors the algorithm behaves better. While for the original algorithm the average PR curve starts below 1 and goes lower as the in-out degree increases, the inclusion of priors and of the new estimates of decay rates leads to always having the top scoring interaction be correctly predicted as true. Of course the slope of the curve increases when the degree of the network increases, but the curves using priors (or priors and decay rates) are systematically above the original ones.

**Figure 7:**
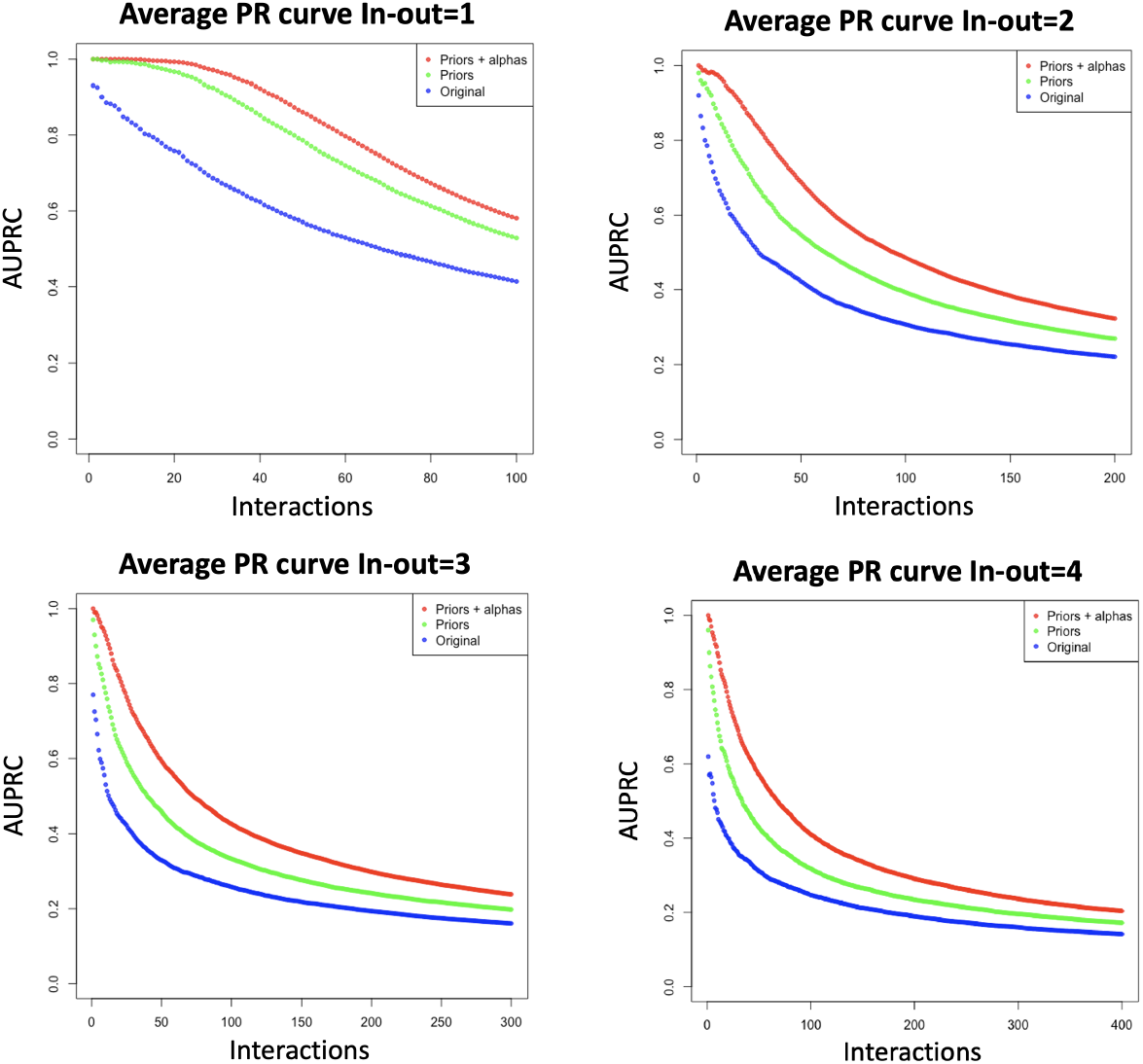
PR curves obtained by averaging over 100 graphs when using the original dynGENIE3 algorithm and our two improvements, for different in-out degrees. The y axis displays the PR values, while the x axis indicates the number of interactions to be assigned as TRUE.

### 3.4 Temperature-based rescaling of priors

In this section, we introduce a useful way to modify our priors. This is done by “reshaping” our (not normalized) priors via a non-linear transformation:

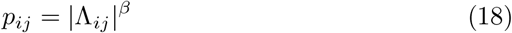

The exponent *β* can be interpreted as an inverse temperature. In the limit *β* → 0 these priors become uniform while for *β* = 1 we recover the original choice.

This transformation is standardly used in Large Language Models (LLMs) and helps LLMs achieve better perfomances [20]. As was shown above, the distribution of time-lagged correlations is very close to Gaussian. Thus the distribution of priors modified by Eq. 18 can be estimated analytically as follows. Starting from:

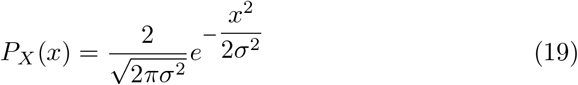

where *x* represents the absolute value of the time-lagged correlation considered as a random variable (we have a factor 2 since we are taking the absolute value of Λ_*ij*_), we define *p* = *x*^*β*^, yielding:

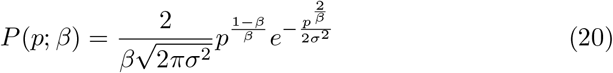

The behaviour of this distribution as a function of *β* is displayed in Fig. 8. We can distinguish two regimes for these distributions: either there is a peak at a strictly positive argument or the peak is at the origin. The position of the peak can be computed analytically from Eq. 20 and we can use it as an “order parameter” to distinguish two regimes of our inference.

**Figure 8:**
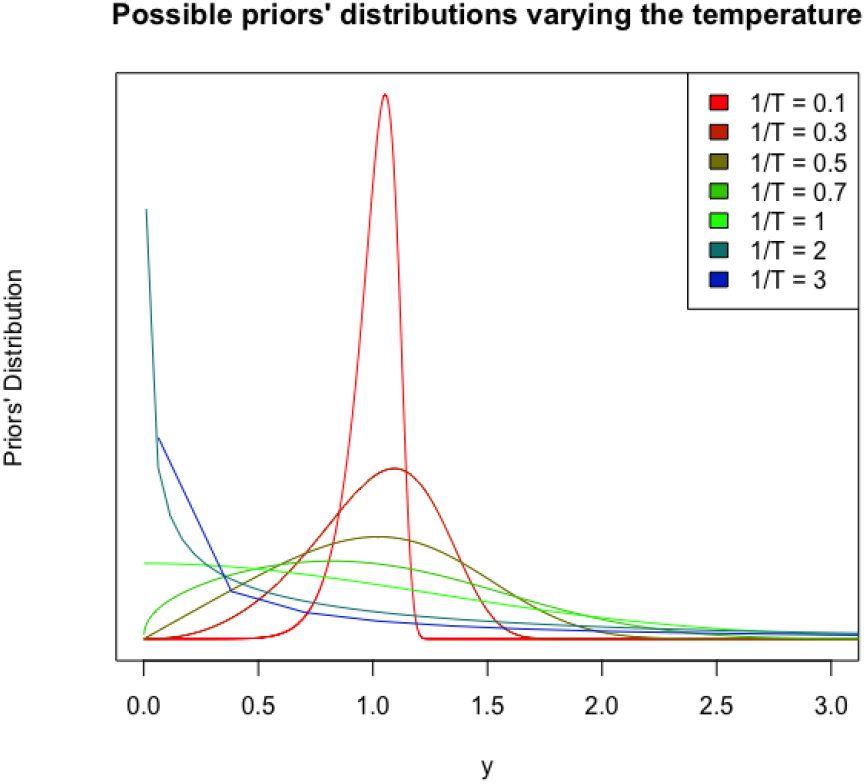
Distribution of the transformed priors at different values of the temperature parameter *T* = 1*/β*, after rescaling Eq. 20 so the means are 1 for all *β*.

The position of the maximum arises at:

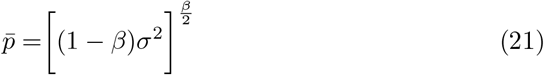

which tells us that *β*_*c*_ = 1 is a critical point. If *β* is much less than 1, corresponding to a “warm” phase, the modified priors will be quite spread out, not distinguishing much the weakly and strongly correlated driver-target pairs. In contrast, when *β* is much larger than 1, corresponding to a cold phase, there is an intense focusing on the highest correlations during the construction of the trees of the random forest. As a result the effective size of the search space will be smaller. To showcase this, we consider the “effective” number of candidate drivers to be considered during the random forest construction.

For a given target gene *j* and its corresponding list of priors *p*_*ij*_, we define its effective number of candidate drivers as:

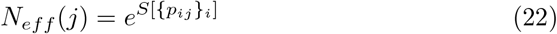

where *S*[ { *p*_*ij*_ }_*i*_] is the Shannon entropy of those priors (at given *j*). In practice, for each in-out degree, we average the *N*_*eff*_ (*j*) over 100 graphs at a given temperature. We then introduce the quantity *N*_*eff*_ as the effective number of candidate interactions obtained by summing all the *N*_*eff*_ (*j*), 1 ≤ *j* ≤ 100. As can be seen in Fig. 9, at *β* = 0, *N*_*eff*_ = 10^4^, that is the total number of possible edges in our system. When we increase *β, N*_*eff*_ decreases for all cases displayed, in line with the intuition that at low temperatures the forest constructions effectively considers fewer drivers. Interestingly, the four curves in Fig. 9 are nearly identical.

**Figure 9:**
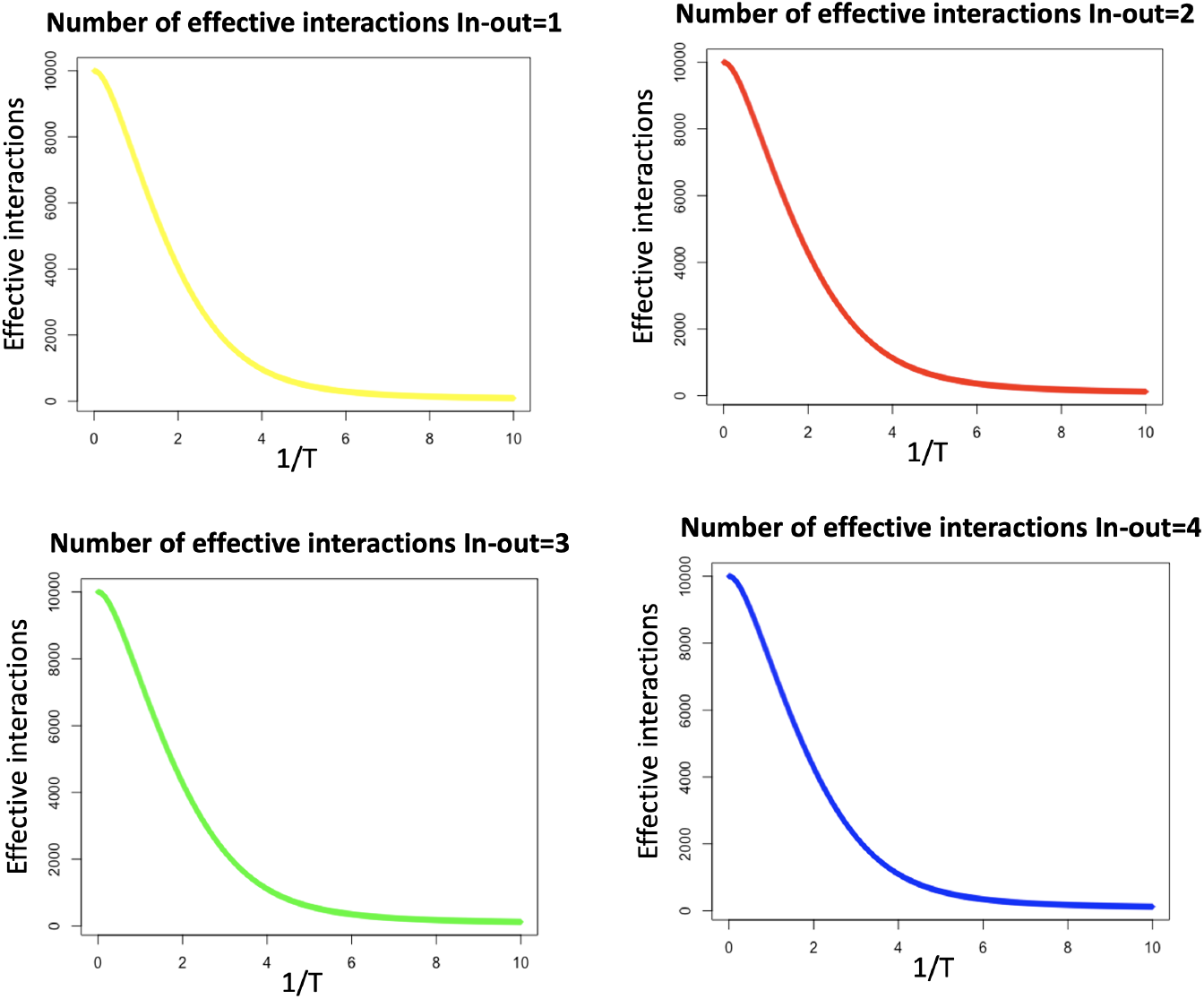
Dependence on (inverse) temperature of the effective number of candidate interactions considered in the random forest constructions, for different in-out degrees. The values are obtained by averaging over 100 random graphs, each having *N* = 100 genes.

Given that temperature allows the random forest search to focus on putatively more relevant drivers, one can expect that temperature can be used to improve the predictive power of dynGENIE3. To test this expectation, we ran our algorithm (including our modified estimates for the decay rates) at different temperatures and computed the AUROC and AUPRC measures of predictive power.

As one can see in Tab. 3 and in Fig. 10, temperature rescaling leads to an increase in the AUPRC, which is higher for smaller in-out degrees, while the AUROC remains mostly unchanged. This can be explained by the fact that the AUPRC measure tends to be sensitive to getting the top scoring interactions right. Such a trend is expected since using more skewed priors by decreasing the temperature focuses the attention on the top candidate drivers as shown earlier. The optimal inverse temperature in this system is around 3 and interestingly it is insensitive to the in-out degree. Note also that this use of temperature works best for low in-out degree, producing only modest improvements for the higher in-out degrees.

**Figure 10:**
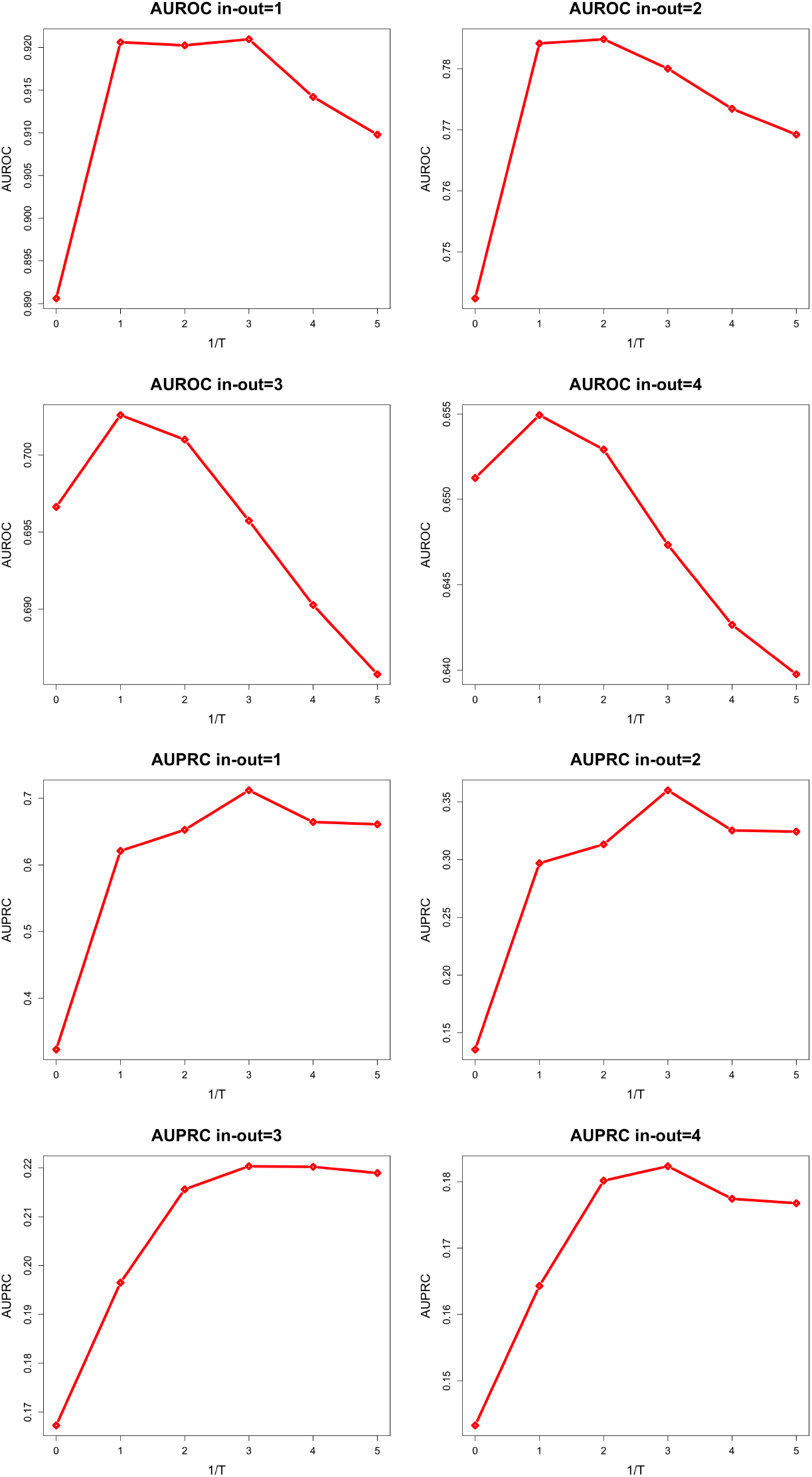
AUROC and AUPRC at different temperatures for different in-out degrees.

## 4 Conclusions

In this paper, we have shown how the inclusion of priors in the dynGENIE3 [14] algorithm can be beneficial for the inference of gene regulatory networks (GRNs), increasing significantly the reliability of predictions. We have focused on using linear correlation coefficients to build our priors, but likely other approaches may also be relevant. We considered both equal-time and time-lagged correlations.

Not surprisingly, we found that the signs of these correlations can be used to determine whether an interaction of the network is inhibitory or excitatory. This constitutes an advance from the original algorithm which provides unsigned weights for each putative interaction as output.

Moreover, we introduced a novel estimate for the mRNA decay rates based on a simple dynamical model. We saw how using both the new estimate and the priors improved the prediction performance. This improvement is particulary impressive for the AUPRC which is doubled, but is also good for the AUROC which is systematically increased. This tells us that by biasing (priorizing) the inference according to linear correlation coefficients, we can improve prediction reliabilty of these random forests approaches at essentially no computational cost ^6^.

It is worth mentioning here that the framework we have developed is general and can be used to include any knowledge about the network. One could, for instance, base priors on biological data, e.g., via ChIP-seq [19] or DAP-seq [6] data that suggest which genes might be regulated by which others. That idea has been explored in other works such as [21]. Our framework is perfectly applicable there since the matrix of priors can include whatever information the user provides.

In the final section, we showed a possible modification of our priors which further improves the performance of the algorithm. Specifically, we introduced a “temperature” parameter to perform a non-linear transformation of our priors. This transformation allowed us to study the effect of enhancing priors on the inference. By changing this temperature, it is possible to follow the inference in different phases and to understand what is the best biasing procedure. In this context we have been able to characterize two phases for the inference in terms of an order parameter, and to establish that the low temperature phase gives better results because it allows the algorithm to focus on a smaller number of candidate interactions.

In conclusion, the present work shows that the inclusion of priors can lead to major benefits in the inference of GRNs. We provided a particular framework and associated computer codes that can predict with much higher accuracy the interactions of a network from time series data. Our result show that with our improvements we are able to double the AUPRC and increase substantially the AUROC. Other avenues of improvement, such as the use of biological information, can be explored within this framework with little effort. It is quite possible that some of the strategies we proposed here in the context of gene regulatory networks may improve random forest inference in more general settings.

## Footnotes

1 Here we only care about the generation of time series data. For information about steady state data, consult [22]

2 This is done as to avoid picking up too small deviations from Gaussian.

3 There are of course exceptions; here we will not worry about more complicated cases as arise for instance when drivers act along with co-factors.

4 Of course one can achieve better performances by increasing the number of drivers to choose from at each split and the number of trees, but this has a computational cost which slows down the inference. Our choice of *n*_*trees*_ = 100 allows for the inference to be quite fast.

5 If we were to expand past the first order we would end up with having to infer the “couplings” between different genes, which are what we want to find once we have estimated the decay rates, so it would be circular.

6 Although we did not test it systematically, running the code with priors can be faster than when there are no priors. Intuitively this may be understood as follows: whenever the distribution of genes to sample is not uniform, the tree construction effectively works with a lower number of candidate interactions and thus will be more efficient computationally.

